# Seq2Neo: a comprehensive pipeline for cancer neoantigen immunogenicity prediction

**DOI:** 10.1101/2022.09.14.507872

**Authors:** Kaixuan Diao, Jing Chen, Tao Wu, Xuan Wang, Guangshuai Wang, Xiaoqin Sun, Xiangyu Zhao, Chenxu Wu, Jinyu Wang, Huizi Yao, Casimiro Gerarduzzi, Xue-Song Liu

## Abstract

Neoantigens derived from somatic DNA alterations are ideal cancer-specific targets. In recent years, the combination therapy of PD-1/PD-L1 blockers and neoantigen vaccines shows clinical efficacy in original PD-1/PD-L1 blocker non-responders. However, not all somatic DNA mutations can result in immunogenicity in cancer cells, and efficient tools for predicting the immunogenicity of neoepitope are still urgently needed. Here we present the Seq2Neo pipeline, which provides a one-stop solution for neoepitope features prediction from raw sequencing data, and neoantigens derived from different types of genome DNA alterations, including point mutations, insertion deletions, and gene fusions are supported. Importantly a convolutional neural networks (CNN) based model has been trained to predict the immunogenicity of neoepitope. And this model shows improved performance compared with currently available tools in immunogenicity prediction in independent datasets. We anticipate that the Seq2Neo pipeline will become a useful tool in prediction of neoantigen immunogenicity and cancer immunotherapy. Seq2Neo is an open-source software under an academic free license (AFL) v3.0 and it is freely available at https://github.com/XSLiuLab/Seq2Neo.

## 1. Introduction

In recent years, immunotherapy, represented by PD-1/PD-L1 blockers, is transforming the treatment of cancer. PD-1 is a protein found on T cells that helps the immune systems in check. The combination of PD-1 and PD-L1 helps keep T cells from killing other cells, including cancer cells, which can result in immune evasion [1–4]. Previous studies have reported that only a small proportion of patients will have a lasting clinical response while most patients will have only a transient response or no response at all [5,6]. The combination of PD-1 blockers and other immunotherapy like neoantigen vaccines presents favorable development prospects [7,8].

Neoantigens derived from somatic DNA alterations are ideal cancer-specific targets. Neoantigen vaccines have demonstrated therapeutic effects in enhancing immunotherapy efficacy [9]. It has also been reported that the combination of PD-1 antibody and neoantigen vaccine is safe and effective for cancer patients [8]. In addition, TCR-T targeting neoantigens have shown dramatic effects in clinics [10]. However, the success of these neoantigen related therapies relies on efficient neoantigen prediction tools.

A plethora of HLA-peptide binding prediction algorithms have been developed to predict which peptides will bind to specific cognate HLA alleles [11–14]. However, HLA-peptide binding affinity alone is not sufficient for predicting the immunogenicity of peptide. In addition, currently available neoantigen prediction tools provide limited neoantigen features or focus on specific genome alterations, such as point mutations. Accurate prediction of the immunogenicity of neoantigen based on raw sequence data is still urgently needed. Here we present an open-source pipeline tool, Seq2Neo, to provide a one-stop service for raw data preprocessing, HLA typing, mutation calling, and neoantigen prediction. Neoantigens derived from point mutations, insertion and deletions (INDELs), and gene fusions are supported. Various neoantigen features including HLA binding affinity, transporter associated with antigen processing (TAP) transport efficiency and gene expression are predicted for each candidate peptide. Importantly, a convolutional neural network (CNN) based immunogenicity prediction model has been constructed, and this model shows improved performance compared with known methods.

## 2. Results and Discussion

### 2.1. Neoantigen features prediction

Seq2Neo uses a command line-based interface, allowing the users to perform the workflows automatically. Seq2Neo uses publicly available tools for mutation calling, HLA typing and HLA affinity binding prediction. Then a CNN-based model was constructed with the features generated with Seq2Neo for predicting the immunogenicity of peptides to stimulate CD8+ T cell response directly. Finally, Seq2Neo outputs various peptide features including immunogenicity score, peptide-HLA binding affinity, TAP transport efficiency and gene expression (Figure 1 and Figure S1). The Seq2Neo (Figure 1) begins with importing raw sequencing data in FASTQ, SAM or BAM format and then parses the user input to select the corresponding workflow to run. Point mutations and INDELs detection can be performed with Mutect2 [15], and gene fusion detection is performed with STAR-Fusion [16]. Subsequently, somatic variant data in VCF format are generated. MHC genotyping is performed with HLA-HD [17]. Before neoantigen prediction, sample somatic variants will be annotated by ANNOVAR [18] or Agfusion [19] to get potential mutant peptides.

**Figure 1.**
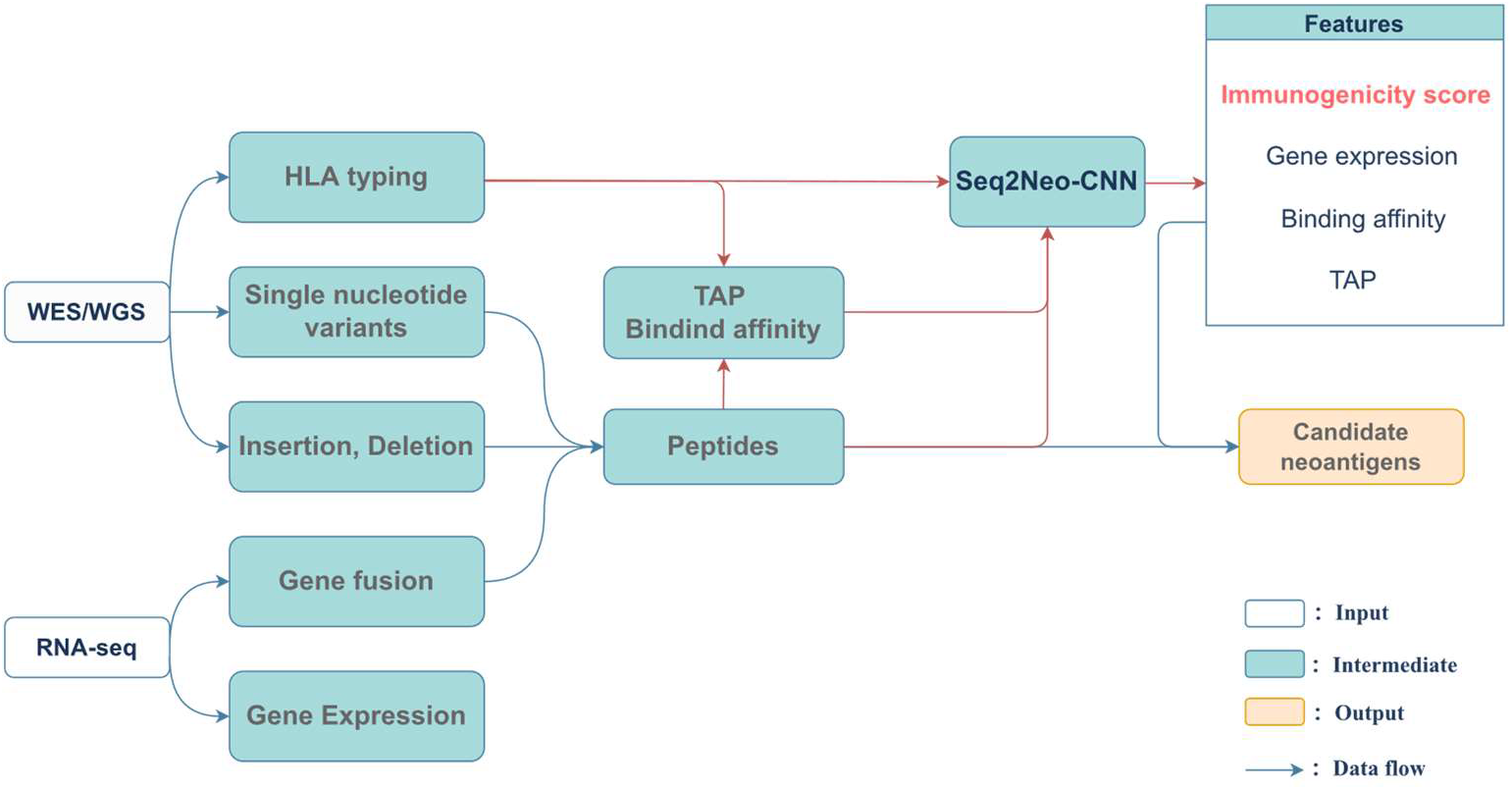
An overview of Seq2Neo. The input of Seq2Neo includes raw WGS, WES, RNA-seq or peptide information. Seq2Neo predicts various peptide features, including a CNN-based immunogenicity score, peptide-HLA binding affinity, TAP transport efficiency, and gene expression. Then Seq2Neo applies those features to rank candidate neoantigens.

### 2.2. Selection of the best HLA-I binding affinity prediction algorithms

We used 23319 peptides (14677 positives, taking IC50=500nm as the threshold) from The Immune Epitope Database (IEDB) to evaluate the performance of peptide-HLA I binding affinity prediction algorithms, including NetMHCpan [20], MHCflurry [21], Pick-Pocket [22], and NetMHCcon [23]. The NetMHCpan BA model obtained the highest accuracy (0.75) and the highest precision (0.96) (Figure 2A-B), and this algorithm was selected for peptide-HLA binding prediction in Seq2Neo. In addition to peptide-HLA binding affinity, Seq2Neo uses TPMcalculator [24] to detect gene expression and obtains TAP transport efficiency from NetCTLpan [25].

**Figure 2.**
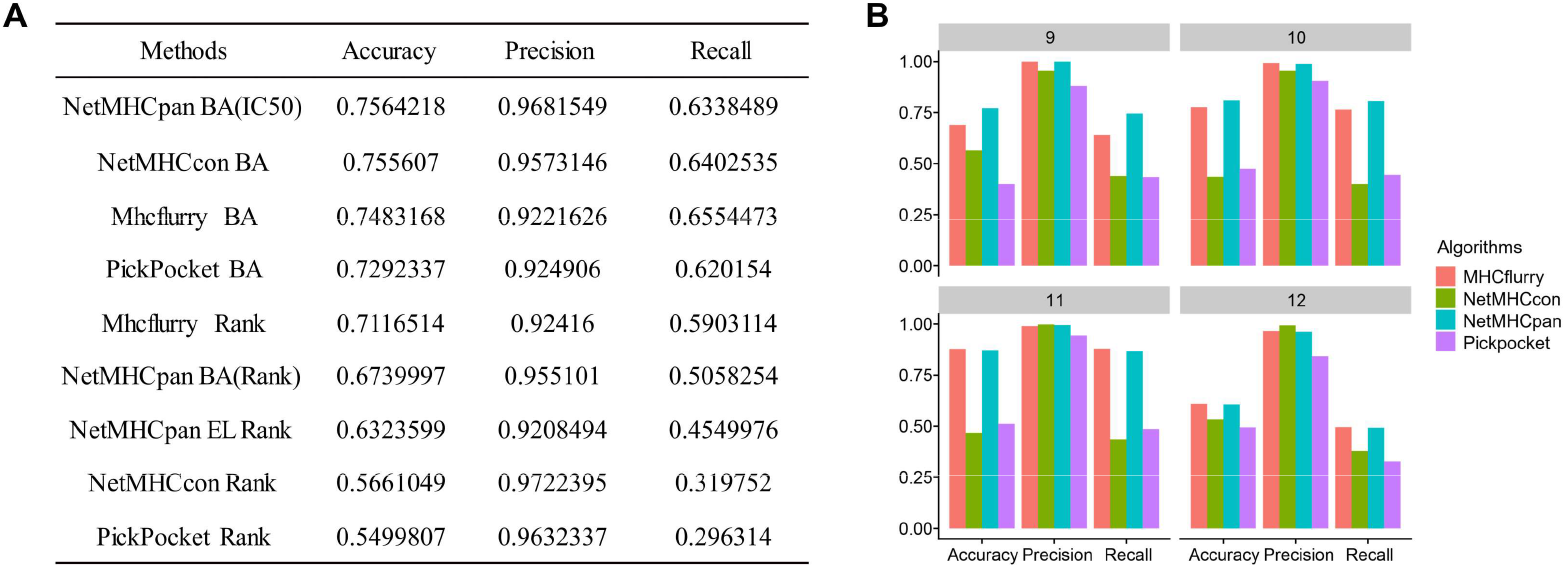
Benchmark analysis of different HLA-I binding affinity prediction algorithms. (A) Comparisons of the performance of different algorithms in prediction accuracy, precision and recall. Dataset was downloaded from IEDB, and the thresholds of IC50 less than 500nM and rank percentile less than 1% were used to determine positive peptides. (B) Comparisons of different algorithms on different lengths of peptides.

### 2.3. Data used for Seq2Neo-CNN model training

The fundamental feature of neoepitope lies in the ability to stimulate cytolytic T cell response, and this immunogenicity information is not predicted with most current neoantigen prediction tools. For immunogenicity prediction, we searched the IEDB database for experimental evidence supporting the immunogenicity of peptides and acquired 75496 experimentally evaluated immunogenicity assays (Figure 3). After applying filter criteria (Materials and Methods), 8975 data instances (5342 negative peptides) were retained in the final dataset. We chose an independent dataset for model validation, with 599 experimentally tested tumor-specific neoantigens from the Tumor Neoantigen Selection Alliance (TESLA) after deduplication and length restriction of 8-11 [26].

**Figure 3.**
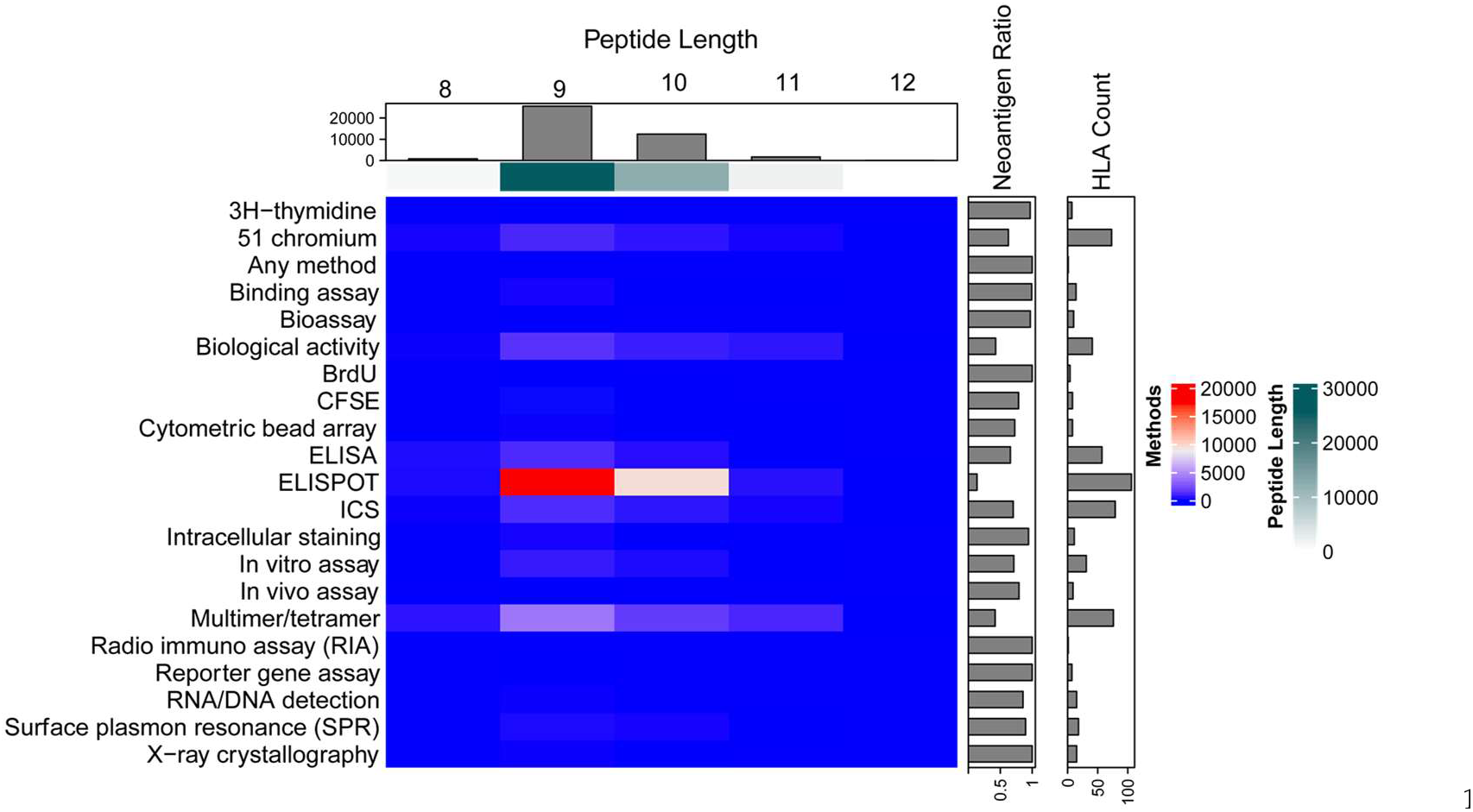
Overview of data used for Seq2Neo model training. Basic information of the IEDB dataset. We restricted the dataset to peptides with metadata that matched the following keywords: (1) linear epitopes, (2) specific T cell assays, (3) intact MHC I class, (4) originated from humans, (5) any diseases, and (6) intact test information for negative peptides. CFSE: Carboxyfluorescein succinimidyl amino ester; ELISA: Enzyme-linked immunosorbent assay; ELISPOT: Enzyme-linked immuno-sorbent spot; ICS: Intracellular cytokine staining.

### 2.4. Features associated with peptide immunogenicity

In order to find some beneficial features for immunogenicity prediction, we compared the features of immunogenic and non-immunogenic peptides. The features of HLA-binding affinity, TAP transport efficiency and proteasomal C terminal cleavage were considered. The differences in HLA-binding affinity and TAP transport efficiency between immunogenic and non-immunogenic peptides were significant but not proteasomal C terminal cleavage (Figure 4A-C). HLA-binding affinity and TAP transport efficiency are not correlated (R = 0.02, P = 0.055, Figure 4D). Therefore, we incorporated these two features into the Seq2Neo-CNN model.

**Figure 4.**
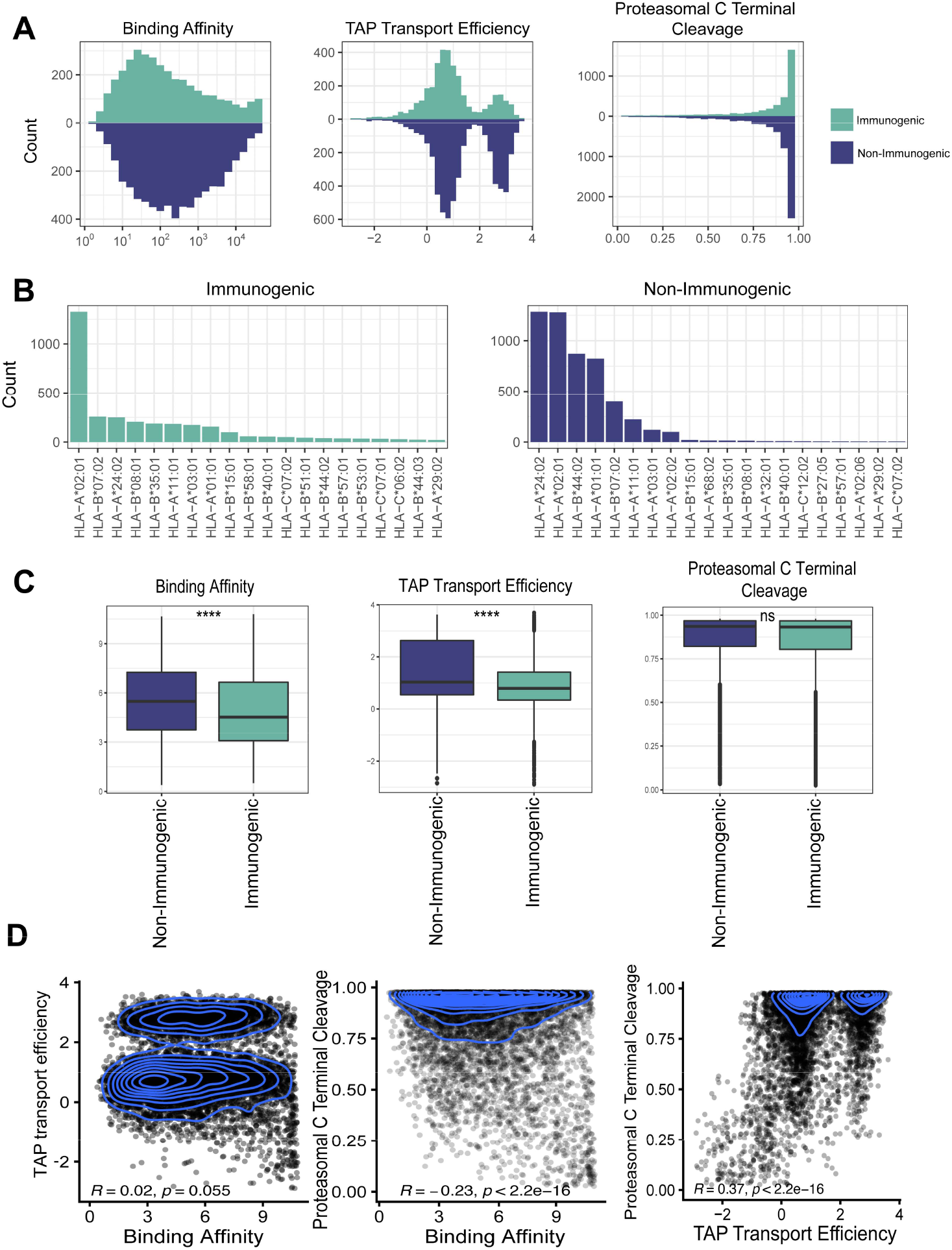
Exploration of the feature differences between immunogenic and non-immunogenic peptides. (A) Distribution comparison of HLA I binding affinity, TAP transport efficiency, and proteasomal C terminal cleavage between immunogenic and non-immunogenic peptides. (B) Distribution comparison of the binding of 20 most frequent HLA-I alleles to the immunogenic (left) and non-immunogenic mutated peptides (right). (C) Comparison of binding affinity, TAP transport efficiency, and proteasomal C terminal cleavage between immunogenic and non-immunogenic mutated peptides. ****: P < 10-4. ns: not significant. (D) Pairwise correlations between the three neoepitope features (peptide-HLA binding affinity, TAP transport efficiency and proteasomal C terminal cleavage). Pearson correlation coefficient R and P values are shown.

### 2.5. Seq2Neo-CNN model for immunogenicity prediction

We built a CNN based model, named Seq2Neo-CNN, to predict peptide immunogenicity (Figure 5A). The performance of the trained CNN model was compared with other machine learning models (ExtraTree, random forest, logistic regression, SVM, and XGBoost) trained with the data collected in this study in terms of prediction accuracy, recall and precision (Figure 5B), and also in independent TESLA dataset (Figure S2). Seq2Neo-CNN model shows the highest performance compared with other machine learning models. The performance of Seq2Neo-CNN was also compared with available neoepitope immunogenicity prediction tools in an independent TESLA dataset, including DeepHLApan [27], IEDB [28], DeepImmuno-CNN [29]. Seq2Neo-CNN also shows the highest performance compared with these known methods (Figure 5C). The details for Seq2Neo-CNN model construction and training are described in Materials and Methods.

**Figure 5.**
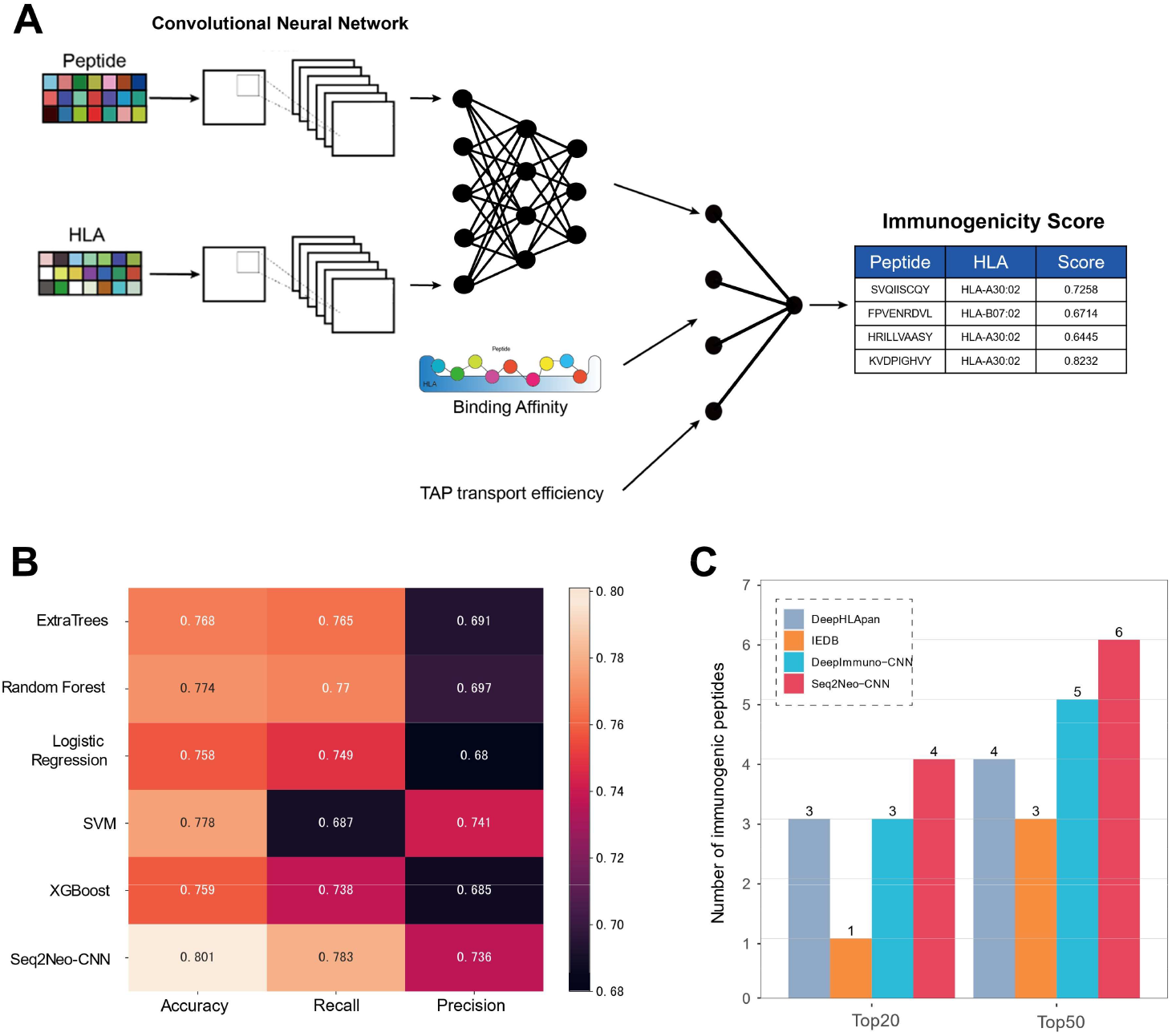
The convolutional neural network-based model (Seq2Neo-CNN) for peptide immunogenicity prediction. (A) The scheme of Seq2Neo-CNN architecture. In this model, each peptide–MHC pair is subjected to two consecutive convolutional layers, followed by three fully connected dense layers, then input to two fully connected dense layers together with TAP transport efficiency and binding affinity information to output an immunogenicity score predictive value. (B) Comparisons of Seq2Neo-CNN with other machine learning algorithms. To select the best predictive model, we constructed five traditional machine learning classifiers (ExtraTree, random forest, logistic regression, SVM, and XGBoost). Each method’s accuracy, recall, and precision were shown. (C) Seq2Neo-CNN outperforms existing immunogenicity prediction methods in the independent TESLA dataset. Comparisons of the performance of different models to predict immunogenic peptides based on the number of true-positive peptides overlapping with each algorithm’s top 20 or 50 predictions.

### 2.6. Seq2Neo validation

In recent years, several tools for predicting neoantigens have been reported. The representative tools of them are shown in the Table 1 [12,14,30–33]. To demonstrate the performance of Seq2Neo, we applied Seq2Neo to five cancer patients with experimentally validated neoantigenic mutations [34–37]. Those cancer samples have WES, RNA-seq data and 16 experimentally validated neoantigenic DNA sites (Figure S3A-B). After applying the selection criteria (TAP >0, IC50 <=500, TPM >0 and immunogenicity >0.5), Seq2Neo identified 10 of 16 validated neoantigenic sites, the rank of the candidate neoantigens was shown in Figure S3C. We then compared the prediction results of Seq2Neo with pVACseq, TSNAD 2.0 and NeoPredPipe, the rank of most validated neoantigenic sites in Seq2Neo was lower than that in the other three, which means Seq2Neo has improved performance in identifying real immunogenic neoantigens (Figure S3C).

**Table 1.** Some representative tools for predicting neoantigens published in recent years. Neoantigen types, input data, neoantigen class, HLA typing, immunogenicity score, TAP score and programming language used are compared.

| Method | Neoantigen types | Input data | Neoantigen class | HLA typing | Immunogenicity score | TAP score | Language | Publish year |
| --- | --- | --- | --- | --- | --- | --- | --- | --- |
| Seq2Neo | SNVs, indels, gene fusions | WES/WGS, RNA-seq | Class I | Yes | Yes | Yes | Python | This study |
| pVACseq | SNVs, indels, gene fusions | VCF | Class I and II | No | No | No | Python | 2019 |
| TSNAD 2 | SNVs, indels, gene fusions | WES/WGS, RNA-seq | Class I | Yes | Yes | No | Python | 2021 |
| NeoPredPipe | SNVs, indels | VCF, HLA types | Class I and II | No | No | No | Python | 2019 |
| Neopepsee | SNVs | VCF, RNA-seq, HLA types | Class I | Yes | Yes | No | Java | 2018 |
| nextNEOpi | SNVs, indels, gene fusions | WES/WGS, RNA-seq | Class I and II | Yes | No | No | Nextflow | 2021 |
| ProTECT | SNVs | WES/WGS, RNA-seq | Class I and II | Yes | No | No | Python | 2018 |

### 2.7. Seq2Neo implementation

Seq2Neo pipeline has been developed with Python 3.7.12 following a clean, modular, and robust design in concordance with best practice coding standards. Instructions on installing and running Seq2Neo are presented in the public GitHub repository (https://github.com/XSLiuLab/Seq2Neo). It has been designed to run as a command line-based program with a user-friendly interface, allowing non-expert users to become quickly familiarized. To facilitate the installation of Seq2Neo, Docker container and Conda package have been provided (Docker: https://hub.docker.com/r/liuxslab/seq2neo, Conda: https://anaconda.org/liuxslab/seq2neo).

## 3. Materials and Methods

### 3.1. Data preprocessing

The Seq2Neo begins with importing data in FASTQ, SAM and BAM format and then parses the user input to select the corresponding workflow to run. The FASTQ files are processed for quality control and removing any adapter sequences at the end of the reads using Fastp [38]. Raw sequence data are aligned to the reference genome (hg38) using the Burrows-Wheeler Alignment tool [39]. If the input format is SAM or BAM, perform GATK best practice first to data preprocessing [40]. The SAM files are sorted, and read group tags are added using the Samtools [40]. After sorting in coordinate order, the BAM files are processed with PICARD MarkDuplicates, and local realignment and quality score recalibration are conducted using the Genome Analysis Toolkit [41].

### 3.2. Somatic mutation detection

Generated or user-inputted co-cleaned BAM files can be used for point mutations and insertion and deletions (INDELs) detection with Mutect2 [15], and gene fusions were detected with STAR-Fusion [16]. Then somatic variant data in VCF format are generated. Besides, parallel computation is enabled, and this significantly reduces computation time.

### 3.3. HLA genotyping

Human leukocyte antigen (HLA) genes play a critical role in antigen presentation and immune signaling. Here HLA-HD [17] is supported for HLA genotyping using DNA-seq data. It will output personal HLA types, including class I and II HLAs for each patient.

### 3.4. Gene expression detection

Being expressed and presented on the surface of tumor antigen-presenting cells is the prerequisite for a neoantigen to be recognized by T cells. Seq2Neo supports the annotation of the expression of neoantigen candidates using TPMCalculator [24].

### 3.5. Neoepitope features

In addition to peptide-HLA binding affinity, other features, including TAP transport efficiency, gene expression, and immunogenicity score can also be predicted with Seq2Neo. These neoepitope features facilitate the filtering of candidate peptides for vaccine or immunotherapy target selection.

### 3.6. Immunogenicity prediction (Seq2Neo-CNN model)

Seq2Neo-CNN as the core of the Seq2Neo pipeline, it can predict the immunogenicity of required peptides. Below, we give a detailed description on the generation of the Seq2Neo-CNN model.

#### 3.6.1. Dataset selection

By using the following IEDB searching conditions: epitope (linear sequence), assay (positive/negative), T cell assay, MHC restriction (MHC class I), host (Human), and disease (any), we collected data from IEDB database for initial training and validation (August 3, 2021 version). In all, we found 75,496 relevant experiments. Although there are different ways to detect the immunogenicity of peptides, some experiments do not detect direct contact with T cells to induce an immune response, so we only selected ELISPOT, 51 Chromium, ICS, Multimer/Tetramer and ELISA validated data. Then, we deleted the instances which do not have 4-digit MHC alleles or are repeated with other instances, limited the length of the peptide to 8 - 11mer and removed negative peptides whose experimental information is missing or tested subjects are less than four. Finally, we got 8975 peptides that meet the requirement in the final dataset, among which 3633 were positive reactive instances and the remaining 5342 were negative. We selected an independent dataset for further evaluation: 599 experimentally tested tumor-specific neoantigens from the Tumor Neoantigen Selection Alliance (TESLA) after selecting 8 - 11mer peptides and removing duplicates [26].

#### 3.6.2. Allele representation

In order to put the MHC class I alleles into the neural network in numerical matrix form, we used the pseudo-sequence to represent it. The pseudo-sequence is constructed by Nielsen et al [42], consisting of amino acid residues in contact with the peptide. The selected positions are 79, 24, 45, 59, 62, 63, 66, 67, 69, 70, 73, 74, 76, 77, 80, 81, 84, 95, 97, 99, 114, 116, 118, 143, 147, 150, 152, 156, 158, 159, 163, 167, 171. We used the following strategy to encode the MHC pseudo-sequences.

#### 3.6.3. Encoding strategy

We used a one-hot encoding scheme to represent each HLA allele and peptide sequence in a numerical matrix, which will be used as input for the following algorithms. The one-hot encoding scheme is realized by assigning a unique integer to each letter in the 21-digit amino acid alphabet containing padding characters as the index of that letter in the amino acid alphabet. Taking the letter “A” for example, we get an alphabet like “ACDEFGHIKLMNPQRSTVWYX” (the unknown amino acid is set to “X”), and the corresponding index of alanine “A” is 0. Then the values of other amino acids are set to 0, but the value of “A” is set to 1. Finally, we get the one-hot vector of [1, 0, 0, 0, 0, 0, 0, 0, 0, 0, 0, 0, 0, 0, 0, 0, 0, 0, 0, 0, 0]. For any peptide, the unique one-hot vectors of each amino acid in its amino acid sequence are vertically combined to form the numerical matrix to complete vectorization.

#### 3.6.4. Features normalization

Binding affinity and TAP transport efficiency are predicted for all peptide-HLAs using the method described previously and then normalized using the maximum and minimum values simultaneously. The basic mathematical form is represented as:

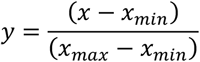

#### 3.6.5. Prediction model

We used CNN (convolutional neural network) to predict the immunogenicity of mutant peptides. The proportion of training set, test set and verification set was 70%, 20% and 10%, respectively. Peptides and MHCs were processed by two consecutive convolutional layers, followed by three dense layers to execute the affine transformation, and then we got a flattened vector of the dimension of 256. NetMHCpan and NetCTLpan were used to calculate the binding affinity (IC50) and TAP of peptides to make use of those as features to train the natural network. To incorporate IC50 and TAP transport efficiency features into CNN, the two dense layers were included. Finally, Seq2Neo outputs the prediction of immunogenicity. We used the ReLu function as the activation function. Some hyperparameters were set through optimizing before starting training, such as batch_size was set to 64, training_loss with patience was set to 15, validation_loss with patience was set to 20, epochs were set to 200 and the learning rate of Adam was set to 0.001. Two early stopping strategies were adopted to ensure that the acquired model was the best. In addition, we adopted batch normalization and dropout strategies to accelerate the model convergence speed and enhance the generalization ability. Since the number of negative reactive instances was significantly more than that of positive reactive instances, the weight was set according to the proportion of negative and positive to eliminate this imbalance. The weight operation is mathematically represented as:

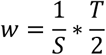

Where w is the negative or positive class weight, S is the number of corresponding reactive instances and T is the total number of training instances.

### 3.7. Other machine learning based immunogenicity prediction models

In order to select the best model to predict immunogenicity, we compared the Seq2Neo-CNN with other five machine learning algorithms (Logistic Regression, SVM, XGBoost, Random Forest and Extra Trees) after optimizing parameters for each method. We used accuracy as evaluation criteria to tune the best parameters for each model. The best parameters of Logistic Regression are acquired through 10-fold cross-validation (penalty=l2, C=2.21). Similar to Logistic Regression, kernel=rbf, gamma=0.1 and C=10 are the best parameters for SVM; max_depth=10, min_child_weight=1.0, gamma=1.625, subsample=1.0 and colsample_bytree=1.0 are the best parameters for XGBoost; n_estimators=200 and min_samples_leaf=2 are the best parameters for Random Forest; n_estimators=1000 and min_samples_leaf=2 are the best parameters for Extra Trees. Then those optimized models were compared with Seq2Neo-CNN in the test set and the TESLA dataset.

### 3.8. Seq2Neo implementation in cancer samples

To test the performance of the Seq2Neo pipeline, WES (normal/tumor exome) and RNA-seq (tumor transcriptome) data from five patients with different solid tumors were downloaded from the NCBI SRA database (bioproject ID: PRJNA298310, PRJNA298330 and PRJNA298376) [34–36]. Each sample has 2–4 experimentally verified neoantigens derived from point mutations which could induce T cell response. Here, we used Seq2Neo to predict those neoantigens for verification of the performance of Seq2Neo. Then we compared the rank percentage of Seq2Neo with that of pVACseq with default parameters.

## 4. Conclusions

As a supplement to PD-1 immunotherapy, neoantigens are ideal cancer-specific targets for precision vaccine design or TCR-T therapy and server as the driving force for cancer immunoediting [43]. However, the current neoantigen prediction steps are cumbersome and lack a comprehensive one-step tool. Furthermore, most neoantigen prediction tools only focus on the binding between peptide and HLA I, and accurate tools for directly predicting the immunogenicity of neoepitope are still lacking. Seq2Neo is a user-friendly and robust tool that provides a one-stop solution for neoantigen prediction from raw sequencing data. Importantly various features of neoantigens can be predicted with Seq2Neo, including the immunogenicity capability of neoepitope.

## Supporting information

Seq2Neo Supplementary file

## Supplementary Materials

Figure S1: The detailed workflow of Seq2Neo; Figure S2: Comparisons of Seq2Neo-CNN and other machine learning algorithms; Figure S3: Application of Seq2Neo in cancer patients.

## Author Contributions

Conceptualization, X.-S.L., K.D. and X.W.; methodology, K.D. and X.W.; software, K.D.; vali-dation, K.D., J.C. and T.W.; formal analysis, K.D., J.C. and T.W.; investigation, K.D., J.C., T.W., X.W., G.W., X.S., X.Z., C.W., J.W., H.Y. and C.G.; resources, X.-S.L.; data curation, K.D., J.C. and T.W.; writing—original draft preparation, K.D., J.C. and X.-S.L.; writing—review and editing, K.D., J.C., T.W. and X.-S.L.; visuali-zation, K.D.; supervision, X.-S.L.; project administration, X.-S.L.; funding acquisition, X.-S.L. All authors have read and agreed to the published version of the manuscript.

## Institutional Review Board Statement

Not applicable.

## Informed Consent Statement

Not applicable.

## Data Availability Statement

HLA-I binding affinity and immunogenicity data were downloaded from IEDB data portal (https://www.iedb.org/, accessed on 3 August 2021). WES data and mRNA expression data from five cancer samples were downloaded from SRA database of NCBI (https://www.ncbi.nlm.nih.gov/sra, accessed on 20 May 2022).

## Acknowledgments

We thank ShanghaiTech University High Performance Computing Public Service Platform for computing services. We thank multi-omics facility, molecular cellular facility of ShanghaiTech University for technical help.

