## Supplementary material for "Seq2Neo: a comprehensive pipeline for cancer neoantigen immunogenicity prediction": Seq2Neo Supplementary file

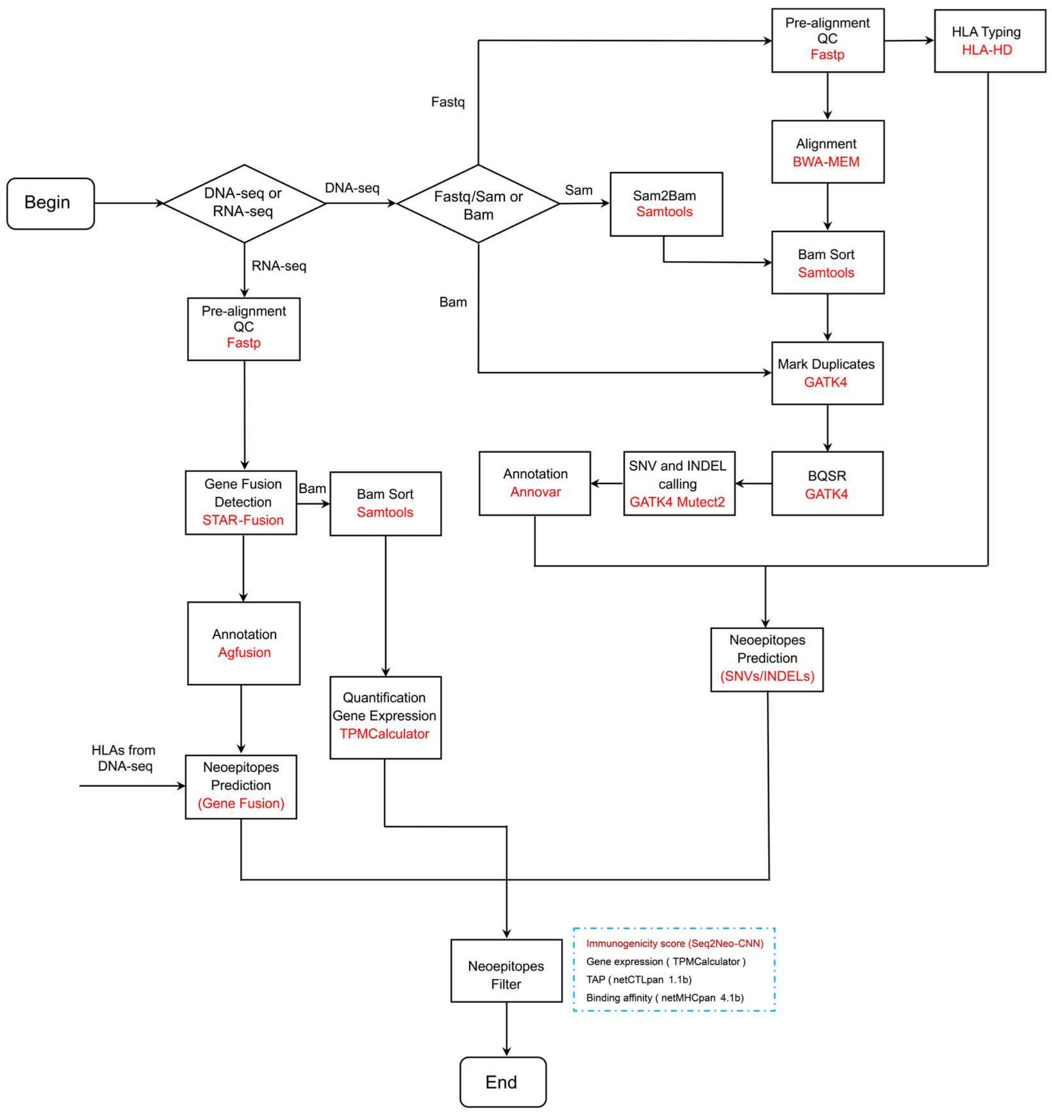


**Figure S1.** The detailed workflow of Seq2Neo. The detailed schematization of the Seq2Neo pipeline, each process and corresponding software are listed.

| Algorithm | Top - 20 | Top - 50 |
|---|---|---|
| ExtraTrees | 1 | 2 |
| Random Forest | 1 | 2 |
| Logistic Regression | 1 | 3 |
| SVM | 3 | 5 |
| XGBoost | 1 | 4 |
| Seq2Neo-CNN | 4 | 6 |

**Figure S2.** Comparisons of Seq2Neo-CNN and other machine learning algorithms. Comparisons of immunogenicity predictions on six machine learning classifiers, with the number of true-positive predictions overlapping each algorithm's top 20 or 50 predictions in the independent TESLA dataset.

A

| Reference genome | Patient ID | #SNVs | #INDELs | #Fusions |
|---|---|---|---|---|
| GRCh38<br>Ensemble release 105 | 3998 | 213 | 14 | 2 |
| | 3942 | 119 | 10 | 4 |
| | 3995 | 59 | 8 | 0 |
| | 4032 | 107 | 12 | 3 |
| | 4166 | 171 | 3 | 1 |

B

| Tumor Type | Patient ID | Validated Mutation Sites |
|---|---|---|
| Melanoma | 3998 | MAGEA6-E168K<br>PDS5A-H1007Y<br>MED13-P1691S |
| Rectal | 3942 | GPD2-E426K<br>NUP98-A359D<br>KARS-D328H |
| Colon | 3995 | RNF213-N1702S<br>TUBGCP2-P265L<br>KRAS-G12D |
| | 4032 | API5-R243Q<br>PHLPP1-G566E<br>RNF10-E572K |
| | 4166 | NPLOC4-I473V<br>SUN1-A127T |

C

| Patient ID | HLA | Peptide | TAP | IC50 | Immuno | TPM | Gene | Change | Rank | Percentage | pVACseq Percentage | TSNAD 2.0 Percentage | NeoPredPipe Percentage |
|---|---|---|---|---|---|---|---|---|---|---|---|---|---|
| 3998 | HLA-A30:02 | SVQIISCQY | 3.131 | 55.08 | 0.7258 | 8.7023 | MED13 | P1691S | 42 | 0.1842 | 0.4706 | NA | 0.2119 |
| 3998 | HLA-A30:02 | KVDPIGHVY | 3.030 | 55.63 | 0.8232 | 4.4051 | MAGEA6 | E168K | 43 | 0.1886 | 0.4853 | 0.4701 | 0.2161 |
| 3998 | HLA-A01:01 | MKVDPIGHVY | 3.147 | 74.73 | 0.6652 | 4.4051 | MAGEA6 | E168K | 57 | 0.2500 | NA | NA | 0.2754 |
| 3998 | HLA-A30:02 | VSVQIISCQY | 3.115 | 76.23 | 0.8130 | 8.7023 | MED13 | P1691S | 58 | 0.2544 | NA | NA | 0.2786 |
| 3942 | HLA-C08:02 | FGDVGSTLF | 1.967 | 15.27 | 0.7359 | 0.5518 | NUP98 | A359D | 7 | 0.0722 | 0.0938 | NA | 0.0596 |
| 3942 | HLA-C08:02 | FGDVGSTL | 0.387 | 68.69 | 0.7433 | 0.5518 | NUP98 | A359D | 23 | 0.2371 | NA | NA | 0.2553 |
| 3942 | HLA-C16:01 | ISKSGLITI | 0.457 | 178.82 | 0.8189 | 0.4029 | GPD2 | E426K | 57 | 0.5876 | 0.5625 | 0.5423 | 0.5574 |
| 3942 | HLA-C08:02 | VVDISKSGL | 0.956 | 248.77 | 0.6974 | 0.4029 | GPD2 | E426K | 66 | 0.6804 | NA | NA | 0.7021 |
| 3995 | HLA-B18:01 | MEHFYLSFY | 2.998 | 8.54 | 0.5300 | 0.3604 | RNF213 | N1702S | 1 | 0.0263 | NA | NA | NA |
| 3995 | HLA-A18:01 | MEHFYLSF | 2.608 | 10.31 | 0.5479 | 0.3604 | RNF213 | N1702S | 2 | 0.0526 | NA | NA | 0.0145 |
| 3995 | HLA-A30:02 | RILLVAASY | 3.292 | 45.59 | 0.6521 | 3.2718 | TUBGCP2 | P256L | 11 | 0.2895 | 0.3000 | 0.1834 | 0.1657 |
| 3995 | HLA-A30:02 | HRILLVAASY | 3.156 | 199.70 | 0.6445 | 3.2718 | TUBGCP2 | P256L | 25 | 0.6579 | NA | 0.7210 | 0.5407 |
| 4032 | HLA-A02:13 | LLQCTQQAV | 0.438 | 26.87 | 0.5670 | 10.5973 | API5 | R243Q | 7 | 0.0432 | NA | 0.0254 | 0.0994 |
| 4032 | HLA-A02:13 | RLLQCTQQAV | 0.457 | 42.37 | 0.5380 | 10.5973 | API5 | R243Q | 11 | 0.0679 | NA | 0.0522 | 0.1551 |
| 4032 | HLA-A03:01 | YLSHLPLTCK | 0.310 | 44.20 | 0.6455 | 10.3660 | RNF10 | E577K | 12 | 0.0741 | NA | NA | 0.1627 |
| 4032 | HLA-A02:13 | ILCETCLIV | 0.413 | 128.13 | 0.6991 | 6.5426 | PHLPP1 | G566E | 65 | 0.4012 | 0.7059 | 0.0501 | 0.4383 |
| 4032 | HLA-B15:53 | RQVILCETCL | 1.484 | 267.47 | 0.7901 | 6.5426 | PHLPP1 | G566E | 107 | 0.6605 | NA | 0.1963 | NA |
| 4032 | HLA-A03:01 | LSHLPLTCK | 0.664 | 317.52 | 0.6905 | 10.3660 | RNF10 | E572K | 121 | 0.7469 | NA | NA | 0.7846 |
| 4166 | HLA-A02:01 | SISQNPFPV | 0.488 | 52.48 | 0.6424 | 6.0091 | NPLOC4 | I473V | 53 | 0.2087 | NA | 0.2182 | 0.2624 |
| 4166 | HLA-B07:02 | FPVENRDVL | 0.488 | 53.43 | 0.6714 | 6.0091 | NPLOC4 | I473V | 55 | 0.2165 | NA | NA | 0.2667 |

**Figure S3.** Application of Seq2Neo in cancer patients. (A) Counts of different types of mutations in the five cancer samples. (B) Experimentally verified neoantigenic sites of the five cancer samples. (C) Predicted neoantigens from the verified neoantigenic sites in the five cancer samples. The last three columns indicate the rank of neoantigens predicted by pVACseq, TSNAD 2.0 and NeoPredPipe, respectively. The lower rank of experimentally verified neoantigenic sites in Seq2Neo compared with other tools suggests the robustness of Seq2Neo in neoantigen prediction.
